# 3D Confinement-enabled Priming of Synaptic Activation Promotes Primary T Cell Expansion

**DOI:** 10.1101/2023.03.02.530690

**Authors:** Ruoyu Jiang, Yu-Hsi Chen, Ritesh Parajuli, Anshu Agrawal, Abraham P. Lee

## Abstract

The success of autologous cell therapy, which depends highly on T lymphocyte expansion efficiency, is often hindered by suboptimal interactions between T-cell receptors and peptide-MHC molecules. Here, we demonstrate 3D confinement-enabled priming of T cell–MHC immune synapse junctions based on cytoskeletal forces within minutes, which is 200-fold faster than conventional 24 h bulk shaking method. Using T cell–Dynabead binding skeletons in the starting culture, two- to six-fold greater T cell expansion was achieved over the conventional T cell expansion approach without inducing excessive cell exhaustion. Under 3D force-confinement, T-cell division (G1, S, and G2 phases) was increased to be twice as fast. Creating 3D T cell–Dynabead skeletons as the “booster” material enables highly efficient T cell expansion, without requiring complex surface modification of antigen-presenting cells. This method can be modularly adapted to existing T cell expansion processes for a wide range of applications including adoptive cell therapies.

**Teaser:** 3D confinement-enabled priming of synaptic activation enables radically faster autologous cell production.

## Introduction

The adaptive immune system depends highly on the clone-specific T cell receptors (TCRs) expressed on the surface of T cells to sense and detect foreign pathogens and malignant cellular changes (1–3). T-cell activation begins with TCRs recognizing antigenic peptides presented by the major histocompatibility complex (MHC) of antigen presenting cells (APCs), and is completed by the presentation of co-stimulation cues (4, 5). Signal transduction to the cytoplasm is initiated upon such binding events; immunoreceptor tyrosine-based activation motif (ITAM) phosphorylation is triggered, followed by a cascade of intracellular signaling events, such as ZAP70 activation and CD4/CD8 receptor-mediated membrane reorganization, that are important for creating stable immunological synapses (6–8). During engagement events between peptide-MHC (pMHC) molecules and TCRs, actin cytoskeleton machinery alters the T cell membrane, restricts the interaction, and concentrates available MHC-peptide complexes by 100-fold to the center of the synapse (9). Active cytoskeletal transport of TCRs exerts a wide range of forces (piconewton to nanonewton) that stabilize immunological synapses; T cell expansion proceeds with the addition of pro-survival cytokines (10–12). The essence of therapeutic adoptive cell transfer (ACT) derives from such T cell antitumor functions, and involves substantial *ex vivo* activation and expansion. In treating melanoma and leukemia, such *ex vivo* expansion of tumor infiltrating lymphocytes and chimeric antigen receptor (CAR) T cells is essential; it requires, at the minimum, billions of T cells for successful clinical tumor detection and eradication (13–16).

Approaches to facilitate *ex vivo* T cell expansion, by stabilizing the robustness of immunological synapses and artificial APCs (aAPCs), have been developed and extensively examined (17–22). One of the most common commercial and clinically relevant approaches is based on Dynabeads functionalized with activating antibodies against CD3 (TCR stimulus) and CD28 (costimulatory cue) (23). Despite the success of polyclonal T cell expansion using these beads systems, the static and rigid bead surface does not fully represent how the APC surface engages with T cells. To address this, various groups have created fluidic lipid membranes coated with ligands for TCRs and costimulatory molecules, mimicking natural engagement of APCs. Fluidic membranes are more effective than rigid spherical beads for creating immune synapse clusters, generating significant T cell expansion (24–27). The high aspect ratio of the bilayer lipid scaffold material (∼70 μm in length and ∼4.5 μm in diameter), or macroporosity of the porous 3D alginate structure, increases their interaction with T cells, resulting in greater proliferation and expansion of human T cells than that of the smaller commercial Dynabeads (4.5 μm diameter), thus providing better anti-tumor activity (28, 29). Exogenous force applied at the piconewton range induces single-cell activation signals (Ca^2+^ flux); this doubles the expansion efficiency when cells and elastic droplet-based aAPCs are cultured under oscillatory shaking conditions (11, 30). Despite advances in the development of novel aAPC materials, the level of complexity involved in creating these materials increases their fabrication time, hampering their mass production for clinical use. Such systems are based on the “bottom-up” hypothesis that randomly distributed T cells and aAPC materials consume energy to engage and stabilize immune synapses, resulting in the formation of T cell–material clusters for clonal expansion (*31, 32*). Further, during the initial interaction stage, the pre-sedimentation T cell–aAPC material contact is randomly governed; this stochasticity places limitations on these emerging methods. The lack of control during the initial contact, and uneven bead-material distribution, therefore leads to suboptimal T cell expansion (*28-30, 33-35*).

We address this challenge by developing a top-down approach, as opposed to the conventional bottom-up T cell expansion strategy. Since the formation of large T cell– material clusters is likely what determines the success of T cell expansion, we speculated that this stochasticity (random cell–aAPC material contact and larger-cluster formation) can be bypassed. By forming T cell–material skeletons during the initial interaction stage to begin the seeding process, we present a highly efficient high-throughput technique, 3D confinement-enabled priming of synaptic activation (3D CPS), which achieves a high contact rate between human T cells and Dynabead skeletons via three-dimensional (3D) confinement. We demonstrate that the high degree of 3D confinement results in highly concentrated and compact mixtures of T cells and Dynabeads, enabling multiple T cells to receive activation cues from multiple Dynabeads; this leads to highly concentrated immune synapse junctions, based on cytoskeleton mechanics (*10, 35–39*). During cell culture, the cell–bead skeletons enhance cell growth, serving as centralized anchor sites to attract more T cells and Dynabeads to grow around them. This method can easily be coupled with a diverse range of APC materials and readily used for the rapid expansion of functional T cells for adoptive cell transfer and other T cell-based cancer immunotherapies.

## Results

### 3D confinement-based priming for T cell activation: Workflow and mathematics

The method to create high contact-ratio T cell–bead skeletons, and the clonal expansion process for seeding, is presented in Fig. 1A. Human T cells are isolated using the negative immunomagnetic approach from whole blood samples of patients with breast cancer. At least 1 million human T cells are purified by removing the human plasma and serum, placed in T cell medium, and concentrated 200–600 fold via serial gravity-based centrifugation, with a final volume of 15 μL. One million Dynabeads are purified and concentrated 5-fold, resulting in a 5 μL bead volume; this is then manually transferred for incubation with the T cell pellets.

**Fig. 1.**
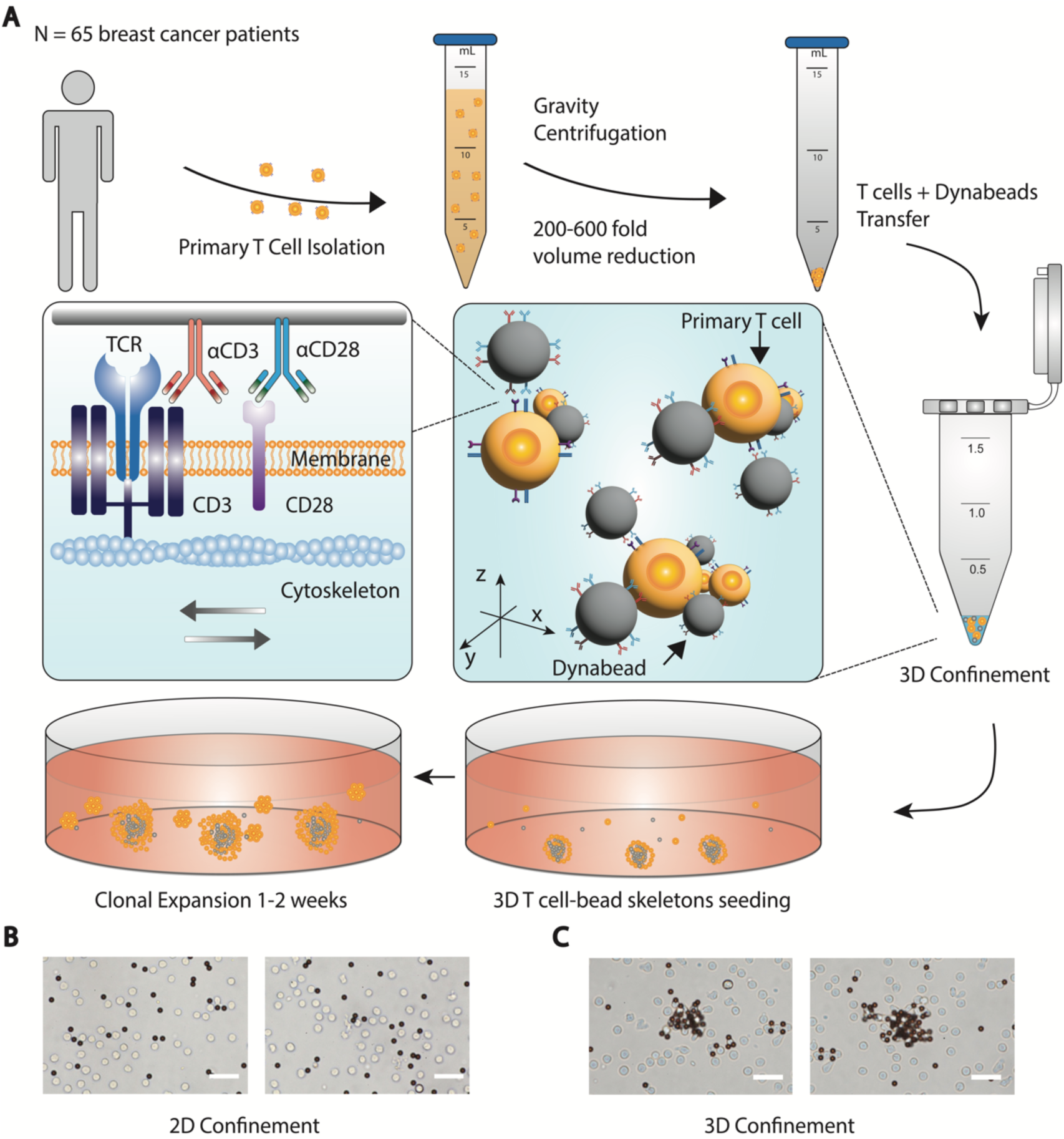
Workflow of 3D confinement-based priming. (**A**). Human T cell pellets are concentrated and confined via serial centrifugation. Stable T cell–bead 3D skeletons are formed from 3D confinement. Seeding is then performed, enabling clonal expansion to continue. (**B**). Representative images of T cells and Dynabeads from 2D confinement. Scale bar: 20 μm. (**C**). Representative images of T cell-Dynabeads skeletons from 3D confinement. Scale bar: 20 μm.

For this purpose, it is important to generate many T cell–bead skeletons very rapidly. The low diffusion speed of T cells and Dynabeads means that few contact events occur before sedimentation. We hypothesized that an extremely high concentration would enable close contact. Interestingly, formation of stable immune synapse junctions was not detected under 2D confinement on flat Countess cell counting slides (Invitrogen) at concentrations of 100 million T cell–Dynabead cluster pairs per mL (T cell:Dynabead ratio, 1:1) (Fig. 1B, Fig. S1A). Most of the interactions were between single T cells and single beads. The lack of physical contact sites prompted us to create a 3D high concentration confinement environment, compacting the T cells and Dynabeads at the bottom of the Eppendorf tube. This ensures that the 3D interaction involves multiple cells and multiple beads, maximizing the contact ratio. The force exerted by the TCR micro-clusters initiates T cell membrane protein reorganization, forming tight immune-synapse junctions (Fig. 1C, Fig. S1B).

Once the cell–bead skeletons are formed in the high concentration environment, the TCR– anti-CD3-antibody bond must be cleaved. Although dense TCR clustering enhances T-cell activation, overstimulation by the TCR stimulus can harm the T cells, causing cell exhaustion (*40*). To address this, 3D confinement was restricted to a short incubation time, then the cell–bead skeletons were transferred to 1 mL cell medium for continued clonal expansion. Since the cell–bead skeletons comprise multiple cells and multiple beads (4–10 T cells and Dynabeads), they are denser than single cells, and thus sediment to the bottom of the well faster. This promotes closer interaction between the cells and beads at the bottom of the well. Further, the T cells expand following activation; this compounding effect increases the cell–bead skeleton size, thus attracting more suspended T cells and beads to participate in the interactions (Fig. S2).

Once T cells bind to Dynabeads, the cytoskeleton actin machinery is initiated, resulting in a series of push and pull movements. The cell–bead skeletons are highly compact and difficult to quantify. The force profiling of single T cells was conducted, assuming that the total force exerted correlates directly with the number of bead–cell interactions. Each oscillation was caused by a protrusive mechanism pushing away from the T cell membrane. The distance travelled by the Dynabead was measured as it was pushed away from the T cell membrane (Fig. 2A). The force generated by the membrane oscillation can be calculated as follows (*41*):

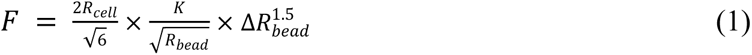

where *R_cell_* is the T cell radius (∼3.5 μm), *K* is the T cell modulus of rigidity, *R_bead_* is the bead radius, and *ΔR_bead_* is the change in bead radius. Commercial Dynabeads are uniformly 4.5 μm in diameter, and the modulus of rigidity of non-activated T cells is 80 Pa (24). Since membrane protrusions were observed from the T cells, the membrane change was calculated as the change in the cell-radius (rather than in the bead radius, as per modifications to Eq. 1 shown in Fig. S3 and Table S1). The traces of the distances were measured, then fitted these using a cubic spine (Fig. 2B). The force generated by this membrane oscillation (Eq. 1) ranges from 3–45 pN (Fig. 2C). The engagement process is rotationally restricted at the interface between the Dynabead and T cell. The polar diagram (Fig. 2D) illustrates that no significant rotation or escape of the bead was observed during the period of tracing. To ensure that this is not caused by Brownian motion, non-interacting beads and cells were also traced and these exhibited no rotational restriction, moving in a random manner (Fig. S4).

**Fig. 2.**
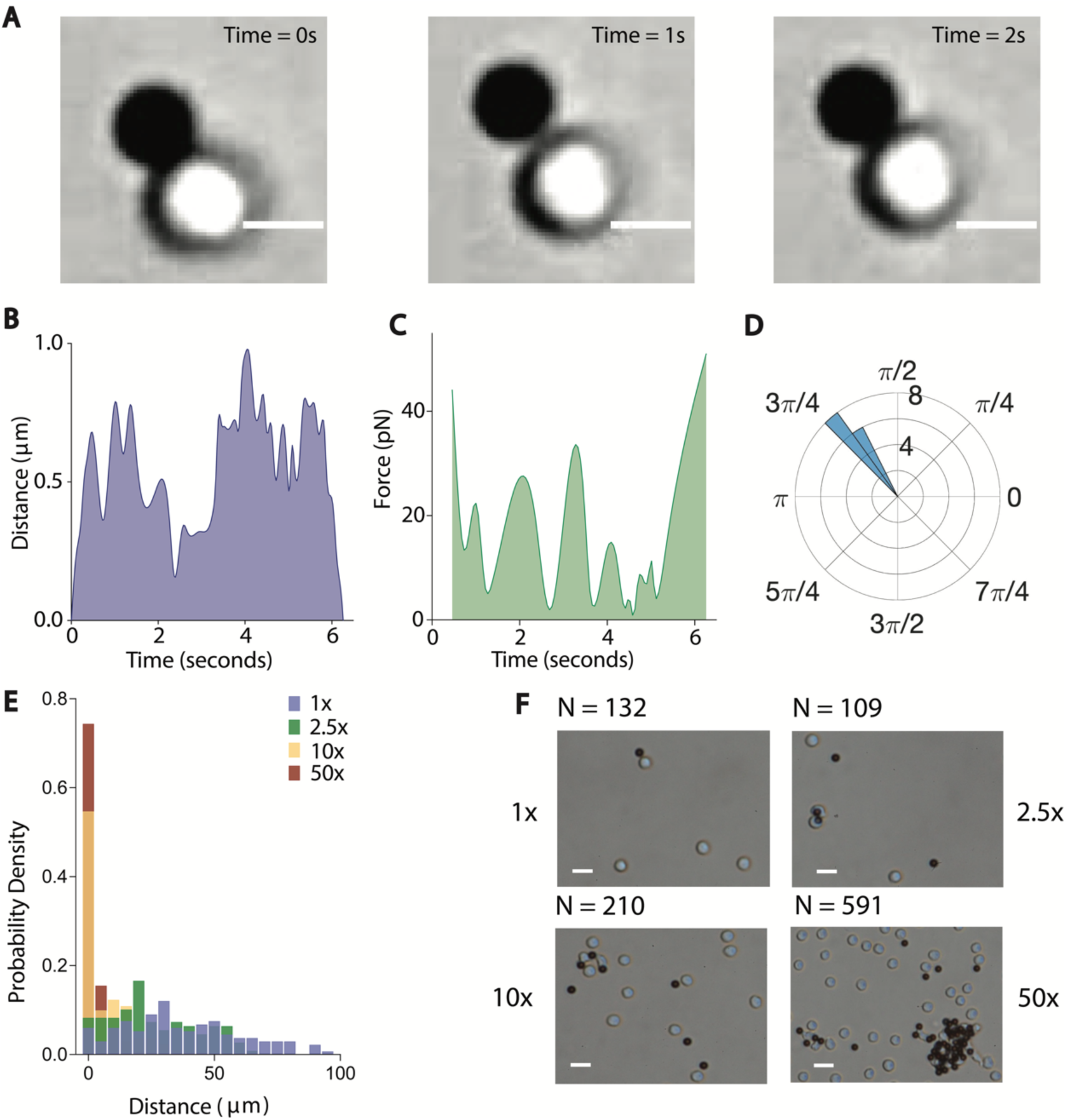
Mathematics of 3D confinement-based priming, and single T cell force profiling. (**A**). Single T cell force oscillatory motion. T cells exhibit substantial oscillatory motion upon interaction with TCR stimuli. Scale bar: 5 μm. (**B**). Distance as a function of time during single cell T cell force profiling. (**C**). The force, resulting from the distance, changes from the edge of the T cells to the Dynabeads. (**D**). Rotational restriction between the T cells and Dynabeads during their engagement period. (**E**). Probability analysis of the close contact distance (μm) under different 3D confinement conditions. (**F**). Representative images for the bulk condition (2 million particles per mL), 2.5-fold concentrated cell solution (5 million particles per mL), 10-fold concentrated solution (20 million particles per mL) and 50-fold concentrated cell solution (100 million particles per mL). N > 100 cells from three independent experiments. Scale bar: 15 μm.

The impact of various dosing concentrations on the interaction behavior were investigated as a function of the cell–bead close-contact (μm) distance. The bulk activation processes were normally distributed, with 12% of the interactions exhibiting a close-contact cell– Dynabead distance peak at 30 μm; only 6% of the cells and beads interacted (Fig. 2E). This distribution restricted the overall interaction rate, suggesting that very few cells were activated during the initial stage. The number of interactions increased as liquid was excluded, and the combined cell and bead concentration increased from 2 million to 100 million particles per mL (Fig. 2, E and F). The distribution peak eventually converged at 0 μm, with 50% of the cell population interacting with beads at 20 million particles per mL, and 80% interacting with beads at 100 million particles per mL. The 40-fold increase in interaction yield achieved in the 3D confinement environment ensures the transport of multiple T cell membrane cytoskeletons, thereby initiating sufficient intracellular activation cascade (Fig. 3A).

**Fig. 3.**
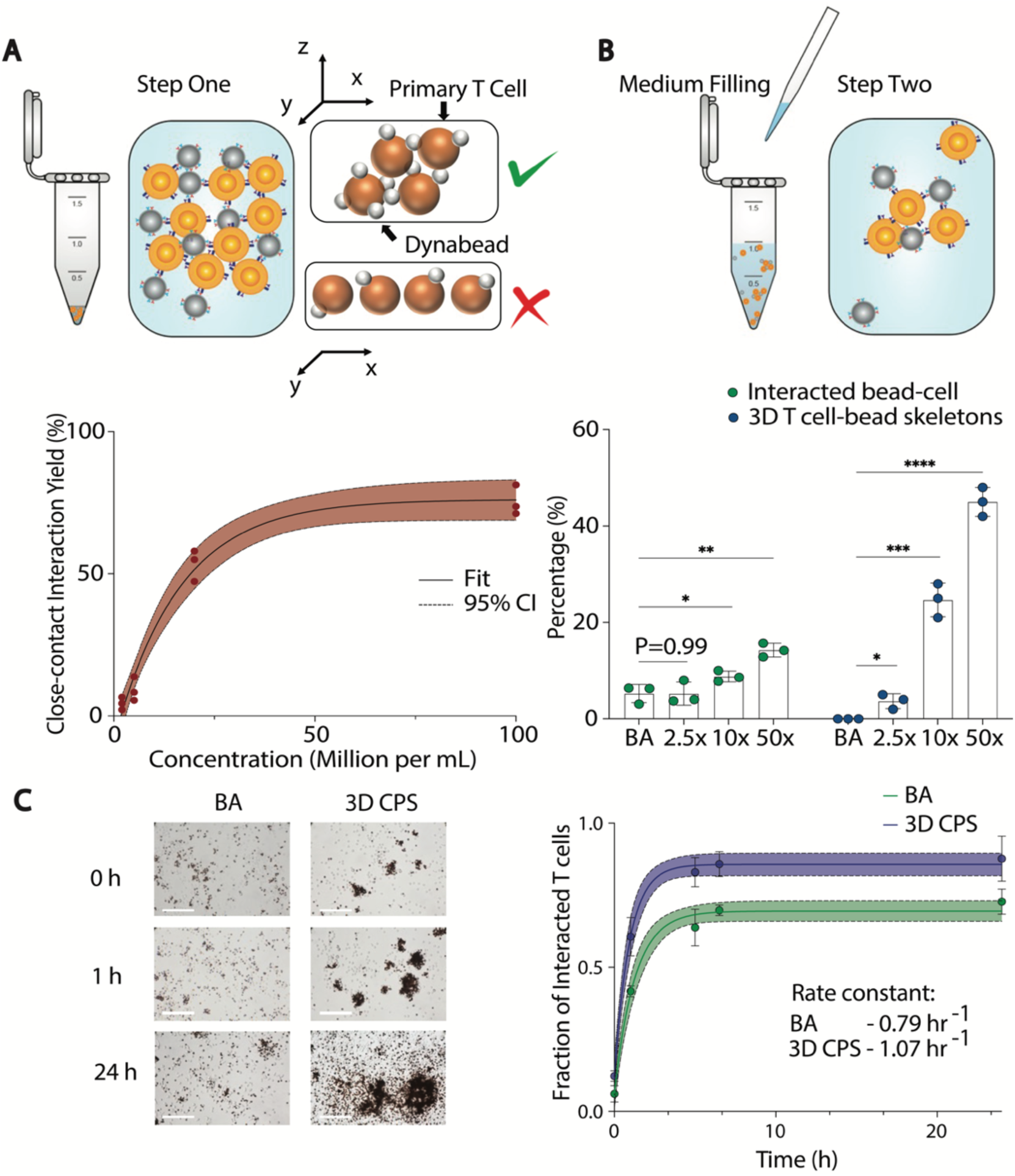
3D confinement-based priming synaptic T cell activation (3D CPS) kinetics. (**A**). One-phase exponential function of 3D confinement concentration (R^2^ = 0.96). (**B**). Seeding process and percentages of cell–bead skeletons. (**C**). Cell–bead skeleton growth over a 24 h period. The fraction of joined cells and beads was used to fit the binding-rate constant. *P* values were determined using unpaired two-tail *t*-tests. * *P* < 0.05, ** *P* < 0.01, *** *P* < 0.001, **** *P* < 0.0001. BA: Bulk activation.

Based on the commercial protocol, the concentration threshold was set at 1 million T cells per mL of medium for culturing purposes (*28, 30*). To avoid overstimulation, culturing of the T cells with Dynabeads under extreme 3D confinement was very brief, and re-seeding of the cells and Dynabeads was performed by transferring them into 1 mL of T-cell medium containing 30 U/mL IL-2, to continue the expansion process. We observed that non-stable immune synapse junctions can be cleaved by manual pipetting, leaving only stable synapse junctions while the conventional bulk process did not generate stable skeletons. In contrast, 45% of the interacted cells formed tight cell–bead skeletons upon seeding (Fig. 3B).

To better understand how the 3D confinement approach affects T cell activation kinetics, the rate of T cell activation was quantified by measuring the fraction interacting during the 24 h culturing process, at concentrations of 2 million and 5 million particles per mL. The interaction status was measured under 40× bright field magnification, imaged the growth process on a 24 well plate, and fitted the fraction of interaction states; the interaction rate constants (*k*) were 0.79 h^−1^ at 2 million particles per mL, and 0.81 h^−1^ at 5 million particles per well (Fig. 3C, Fig. S5, S6). The interacted fractions for these two concentrations remained at 73 ± 4.4% and 71 ± 4.8% at 24 h, suggesting that nearly a quarter of the seeded cells did not engage with the Dynabeads. At 20 million and 100 million per mL, the rate constants were 0.95 h^−1^ and 1.07 h^−1^, respectively; at 24 h, their interacted fractions were 88 ± 11% and 87 ± 7.9%, respectively, suggesting that most of the cells engaged with the Dynabeads for activation (Fig. 3C, Fig. S5, S6). This higher activation rate constant is primarily due to the greater number of cell–bead skeletons present in the starting materials; these skeletons serve as large scaffolds to enhance cell–bead adhesion.

### Polyclonal activation of primary T cells from breast cancer patients

To assess the activation responses, the phenotype of the cultured human T cells was evaluated. The 3D confinement group consistently showed much larger cell–bead clusters than the bulk activation group. Human T cells that were not incubated with Dynabeads exhibited no cluster formation (Fig. 4A, Fig. S7). The 3D confinement approach achieved more centralized clusters, whereas the bulk approach generated scattered T cell clusters. The levels of activation marker CD25 were analyzed on day one, to examine activation efficiency: the 3D confinement approach generated much higher CD25 expression than the bulk approach (Fig. S8). Interestingly, CD4 expression was also significantly higher for the 3D confinement approach (Fig. 4B, C). This is consistent with findings from other studies, suggesting the importance of CD4 in the activation cascade in human cells (*30*). Because CD25 is also used to identify T regulatory cells, CD69 expression was analyzed to verify the activation signal: CD69 expression was similarly elevated following the 3D confinement approach (Fig. 4D).

**Fig. 4.**
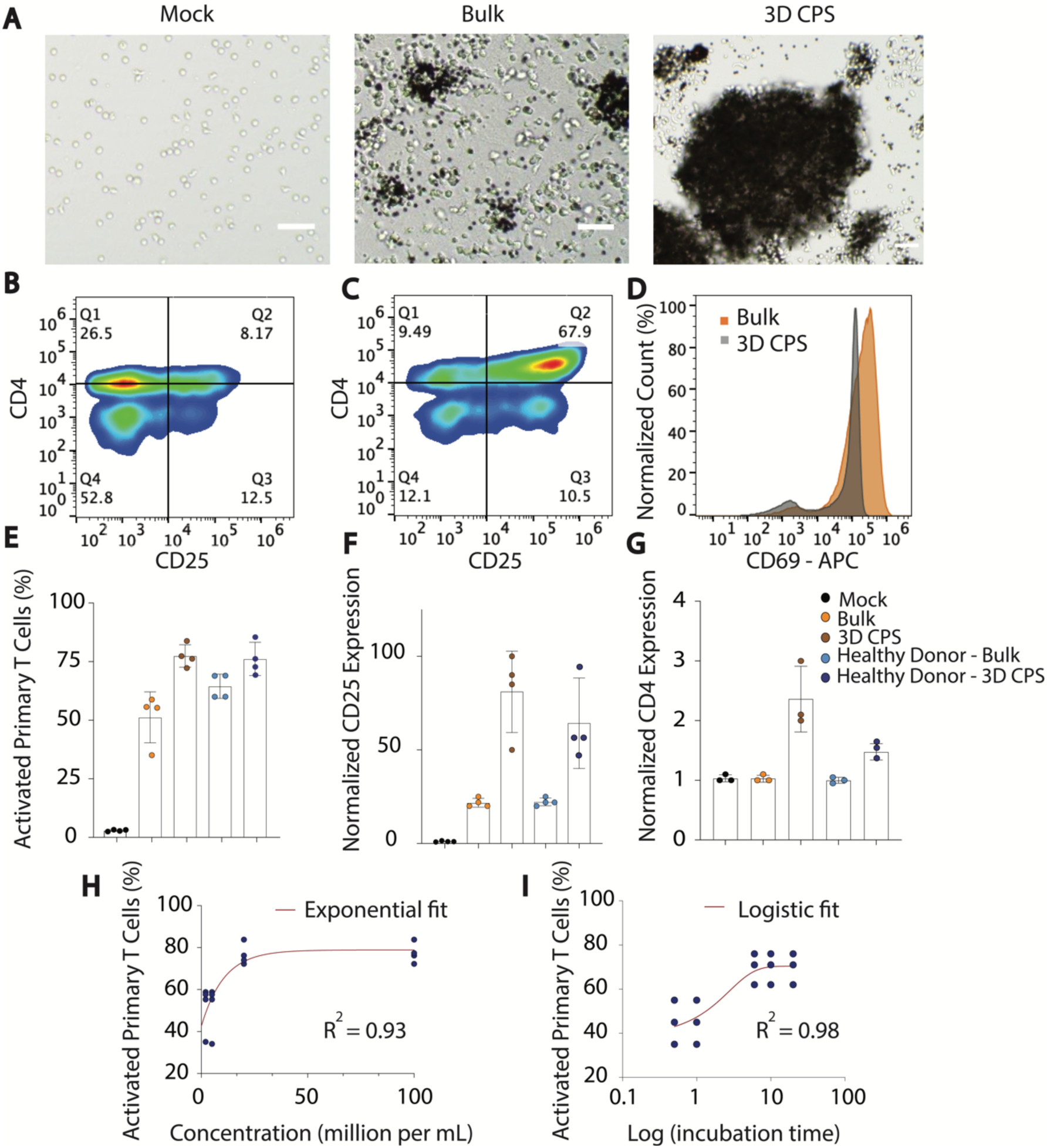
Polyclonal activation of primary T cells. (**A**). Representative bright field images of activation phenotypes for the mock (without Dynabeads), bulk (with Dynabeads), and 3D confinement groups. Scale bar: 50 μm. (**B**). Activation marker CD4 and CD25 analysis for bulk activation. (**C**). Activation marker CD4 and CD25 analysis for 3D confinement groups. (**D**). Activation marker CD69 analysis for bulk and 3D confinement activation. (**E**). Percentage of activated human T cells. (**F**). Normalized CD25 MFI expression. G. Normalized CD4 MFI expression. *P* values were determined using unpaired two-tail *t*-tests. * *P* < 0.05, ** *P* < 0.01. (**H**). Percentage of activated T cells as a function of confinement concentration. (**I**). Percentage of activated T cells as a function of incubation time of 3D confinement at 100 million particles per mL.

Using 3D confinement, the activated T cell fraction increased from 50% to 75% for cancer-patient T cells, and from 60% to 75% for healthy donor T cells (Fig. 4E). For CD25, mean fluorescence intensity (MFI, and indicator of activation expression) was on average three-fold greater for cells from cancer patients, and two-fold greater for those from healthy donors, after 3D confinement relative to the levels achieved by bulk activation (Fig. 4F). For CD4, MFI was on average two-fold greater for cells from cancer patients, and 1.5-fold greater for those from healthy donors, after 3D confinement relative to the levels achieved by bulk activation CD4 (Fig. 4G). The slightly lower signal elevation after 3D confinement in the cells from health donors may reflect intrinsic differences between primary T cells from cancer and healthy donors, possibly due to the antigen-exposure levels in the tumor microenvironment prior to the T cell harvesting (*42, 43*).

CD25 expression was higher at the two higher particle confinement concentrations (20 million and 100 million particles per mL) than at the two lower concentrations (2 million and 5 million particles per mL) (Fig. 4H). To achieve the higher activation signal via the 3D confinement approach, a time threshold of 3–6 min was required (Fig. 4I), consistent with the time needed to form the cell–bead skeletons. At times shorter than this 3D confinement incubation time, T cell activation was not enhanced. Notably, for both the 3D confinement and bulk groups, no CD25 expression was observed during the remaining 23 h when the beads were removed after 1 h of culturing. This suggests that the duration allocated for immune synapse formation is highly important for successful T cell activation.

### Polyclonal expansion of primary T cells from breast cancer patients

The polyclonal expansion of primary T cells from breast cancer patients was investigated by culturing the cells for a two-week period and observing expansion on days 3, 7, and 14. The higher activation signal using the 3D confinement approach consistently generated larger T cell–bead clusters; the bulk expansion group had much lower expansion rates and generated smaller T cell–bead clusters throughout the two-week period (Fig. 5A). This was supported by the much larger T cells observed for the 3D confinement group than for the bulk group; the size distributions showed larger cell diameters on day 3, 7 and 14 (Fig. S9).

**Fig. 5.**
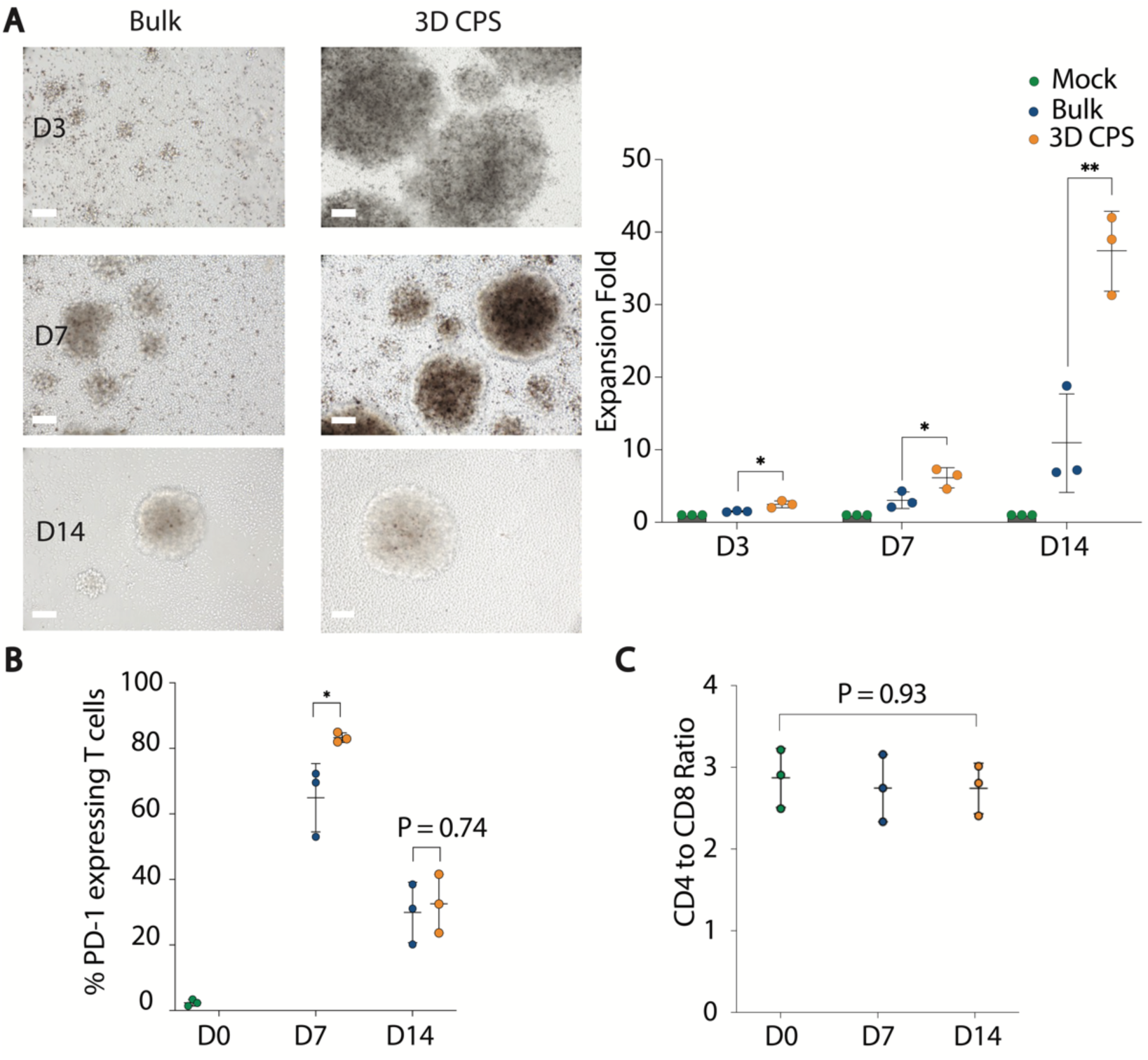
Polyclonal expansion of primary T cells. (**A**). Representative bright field images of expansion phenotypes and expansion fold for the bulk (with Dynabeads) and 3D confinement groups (100 million particles per mL) on days 3, 7, and 14. Scale bar: 50 μm. *P* values were determined using unpaired two-tail *t*-tests. * *P* < 0.05, ** *P* < 0.01. (**B**). Quantification of T cell PD-1 expression over the two-weeks of clonal expansion. (**C**). CD4:CD8 ratio among the CD3^+^ cells on days 0 and 14 of T cell expansion.

Supported by the enhanced T cell activation, programmed cell death protein 1 (PD-1) expression was significantly higher in the 3D confinement group than in the bulk expansion group on day 7, decreasing gradually to a similar level as the bulk group; this suggests that the 3D confinement approach did not induce severe cell exhaustion (Fig. 5b, Fig. S10). Although CD4 expression was much higher on day 1 in the 3D confinement group than in the bulk expansion group, no significant CD4 T cell expansion was observed, and the CD4:CD8 ratio was 2:1 on days 7 and 14, similar to the ratio prior to activation (Fig. 5C, Fig. S11).

The proliferation of the expanded T cells was examined using flow cytometry to evaluate the expression of ki-67, an intracellular marker used to validate the proliferation index and expressed only in T cells undergoing cell mitosis cycles (G1, S and G2 phases). Ki-67 expression increased significantly (by 80.87%) under the 3D confinement approach, by 11.99% under the bulk approach, and by 0.12% for the non-activated CD3^+^ cells (Fig. 6A). Although higher ki-67 expression was associated with higher T cell proliferation, it did not directly reveal which cell division phases the T cells were undergoing.

**Fig. 6.**
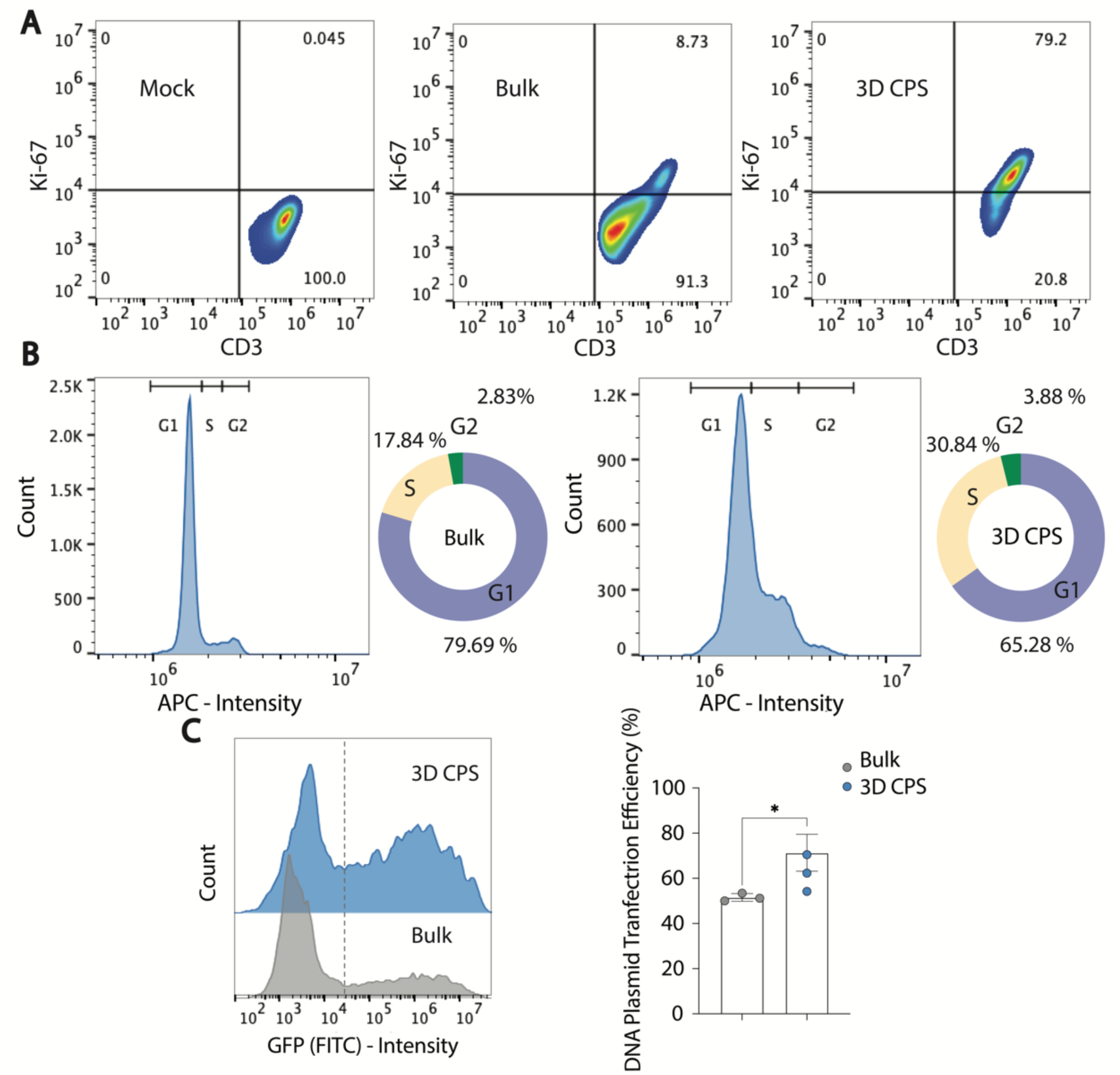
Proliferation of primary T cells. (**A**). Intracellular ki-67 expression under the different expansion approaches. (**B**). DNA mass cycles. (**C**). DNA plasmid transfection (eGFP) efficiency using the bulk and 3D confinement approaches. *P* values were determined using unpaired two-tail *t*-tests. * *P* < 0.05.

Analysis of DNA mass via flow cytometry clearly revealed that the 3D confinement approach enabled the T cells to transition toward the S and G2 phases (related to DNA replication and cell division); these T cells were therefore undergoing rapid cell membrane and nucleus expansion. In contrast, under the bulk approach, the T cells remained primarily at the G1 phase (cell growth) (Fig. 6B).

To further validate this, Lonza electroporation was conducted on cultured primary T cells, which were transfected with the green fluorescence protein (GFP) DNA plasmid. Deformation of the cell membrane and nucleus enhances plasmid uptake by the nucleus. Following electroporation, plasmid transfection efficiency was significantly higher for T cells using the 3D confinement approach than those using bulk approach protocol (Fig. 6C, Figs S12, S13).

## Discussion

3D CPS is a straightforward approach that can be readily adopted for primary T cell activation and expansion, with broad applicability in cell biology, immunology, and development of cell immunotherapy protocols. Since we have demonstrated 3D CPS using commercially available standard Dynabeads, it can be applied by general labs and clinics for research or biomanufacturing. The method generates high contact-ratio T cell–bead skeletons within minutes, as opposed to the 6–8 h required by current protocols, establishing it as a highly efficient approach for robust immune synapse formation. The long formation time required in standard protocols is due to their lack of an architectural design strategy to increase interactions during the initial culturing period before cell sedimentation. Current T cell expansion research focuses on engineering immune synapses after sedimentation, often using aAPC materials, a more bottom-up approach (Fig. S14). Because 3D CPS provides such substantially faster reaction times within small reaction volumes, it has potential for ultra-high throughput: each run (or tube) can accommodate up to 10 million T cells, and a centrifuge can accommodate up to 10 tubes with basic serial passaging. Further, Dynabeads provide a static and rigid aAPC material that can be integrated with fluid-lipid bilayer membrane-aAPC materials, to mimic how T cells engage naturally with APCs.

Polyclonal T cell expansion can be extended to antigen-specific T cell expansion, by coupling the aAPC material with relevant tumor-antigen peptides, or expanding CAR-T cells. Here, we have demonstrated the method using breast cancer patient-derived T cells. Therefore, 3D CPS may also be useful in autologous monocyte-derived dendritic cell (moDC) therapy in clinics, to induce antigen-specific expansion of therapeutic T cells (*44–46*). To enable its widespread application in allogeneic cell therapy, we established that 3D CPS achieves healthy donor T-cell polyclonal activation: CD25, CD69, and CD4 all exhibited elevated activation. Proliferating cells expand their nuclear envelope during mitosis (the M phase) to allow more effective entry of DNA into the nucleus. This expansion during cell division triggers physical enlargement, causing protrusions to extend from the cellular membranes and from the nucleus. The significant elevation of GFP expression following transfection suggests that 3D CPS could contribute to the rapidly growing field of CRISPR-based clinical trials, including those examining targeted knockout of immunosuppressive factors such as PD-1, of endogenous TCR and MHC-1, or studies examining CAR insertion among autologous and allogeneic T cells (*47–49*). Critically, our technique maximizes the surface density cues presented to the T cells without inducing cellular exhaustion. T cells become overstimulated when they are exposed to excessive stimulatory molecules for long periods of expansion, such as weeks. 3D CPS bypasses this obstacle by forming high contact-ratio skeletons as scaffolds before expansion, while maintaining the recommended culture concentration ratio (1:1 for Dynabeads:T cells). Our findings reveal similar levels of exhaustion (reflected by elevated PD-1 expression) in both the bulk activation and 3D CPS groups, with 3D CPS inducing more extensive T cell expansion.

3D CPS, inspired by a bottom-up/top-down development approach, represents a new paradigm for cell expansion. In the longer term, this method has critical advantages over conventional cell expansion approaches, not just in T cell-based therapy, but also in the rapidly growing area of antibody engineering. The critical step in 3D CPS that creates the T cell–bead skeletons can also be applied to T cell-dependent B-cell stimulation. B cell expansion requires robust interactions between B cell receptors and other free soluble antigens, and between T helper cells and B cells via CD40 and MHC-II ligand binding. Rapid B cell clonal expansion forms B cell–T helper cell clusters; subsequent differentiation causes the maturation of B cells, which then produce antibodies such as IgG, IgA, or IgE (*50–54*). 3D CPS therefore has the potential for large-scale antibody generation and related antibody selection in drug and vaccine development.

In conclusion, 3D CPS addresses a fundamental cell–material interaction challenge, by forming T cell-Dynabead skeletons via 3D confinement prior to culturing. In contrast, traditional T cell expansion relies on surface engineering of materials to regulate interactions. We demonstrated a 200-fold improvement over traditional bulk stimulation to enhance T cell–biomaterial interactions. Using standard Dynabeads, this method induced substantial T cell expansion, without inducing notable cell exhaustion.

Nonetheless, our method has potential limitations. For instance, the T cell–bead skeletons contain multiple T cells and multiple Dynabeads within highly concentrated scaffolds. It is therefore possible that multiple Dynabeads could engage with a single T cell, resulting in overstimulation of a small proportion of T cells. This overstimulation was likely not revealed by our bulk protein analysis. In addition, although we focused primarily on polyclonal T cell activation and expansion, it is likely that the TCR binding and activation pathways depend on the T cell subtype; further, the associated TCRs have various binding affinities toward an array of antigens (*55, 56*). To address this limitation, the T cell subtypes can first be sorted, and the Dynabeads can be modified to incorporate various pathogenic peptides, prior to activation and expansion. Last, we were unable to measure the rate of membrane protrusion generated by the TCRs toward the Dynabeads under 3D confinement. High resolution methods to image and measure these forces may be helpful to understand the reported time and concentration thresholds.

In future, it would be interesting to investigate whether soluble antibodies also induce skeleton formation under extreme 3D confinement. We plan to study how this method can be adapted for use with tumor infiltrating lymphocytes, and to examine its anti-tumor activity via co-culture with cancer cells or solid tumors in a mice model. This fundamental advance in immunology—the generation of stable immune synapse junctions via 3D confinement, within minutes—is applicable in a broad range of aAPC material-based systems, offering a simple way to radically accelerate and enhance T cell activation and expansion.

## Material and Methods

### Reagents and cell culture

Primary human T cells were cultured in T-cell media supplemented with 30 U/mL recombinant human IL-2, unless otherwise stated. T cell culture medium was obtained from Gibco RPMI 1640 (Thermo Fisher Scientific, Waltham, MA) supplemented with 10% HI-FBS, 2 mM L-glutamine, 1 mM sodium pyruvate, 50 μM beta-mercaptoethanol, 0.1 mM non-essential amino acids, 1 mM sodium pyruvate, 10 mM HEPES, and 1% penicillin–streptomycin. Human CD3/CD28 T-cell expansion Dynabeads were purchased from Thermo Fisher Scientific.

### Primary T cells isolation

Primary human T cells were obtained from breast cancer patients who had previously consented to participate in a Univerisity of California, Irvine (UCI) IRB-approved clinical protocol permitting blood collection (Clinical Trial UCI-17-43). Total T cells were purified using negative immunomagnetic kits (STEMCELL Technologies, Vancouver, Canada).

### Primary T cell immunophenotyping

Step-by-step protocol was followed for flow cytometry analysis for cell suspension samples (Biolegend, San Diego, CA). For T cell activation experiments, the cell suspensions were stained concurrently with anti-CD25 and anti-CD69 antibodies. For T cell expansion experiments, the cell suspensions were stained concurrently with anti-PD-1, anti-CD4, and anti-CD8 antibodies. The samples were then washed twice with PBS+ by centrifugation, and analyzed on a NovoCyte 3000 Flow Cytometer (ACEA Biosciences, San Diego, CA). Flow cytometry data were compensated using stained single-cell samples. Gates encompassing the positive and negative subpopulations within each compensation sample were used to calculate a compensation matrix in FlowJo (FlowJo, Ashland, OR).

### Polyclonal Primary T cell expansion studies

T cells were enumerated with a hemocytometer using Trypan blue exclusion. Expansion fold was calculated as follows: (The number of total live cells at the respective time point) / (the number of live cells seeded at the start of culture).

### Intracellular ki-67 staining

Patient-derived primary T cell were expanded with Dynabeads for 3 d. The cells were then washed twice with PBS and centrifuged at 350 *×g* for 5 min. Then, 3 mL of cold 70% ethanol (−20 °C) was added to the cell pellets drop by drop, followed by gentle vortexing and incubation at −20 °C for 1 h. The cells were then washed twice with PBS and resuspended at 1 × 10^6^ cells per mL. Then, 100 μL of cell suspension was mixed with Pacific Blue anti-human Ki-67 (Biolegend), followed by incubation at room temperature in the dark for 30 min. The cells were then washed and resuspended in PBS for flow cytometry.

### Cell cycle analysis

Vybrant DyeCycle Ruby stain (Thermo Fisher Scientific) was equilibrated to room temperature. On day 3 of expansion, 0.5 mL of T cell suspension was prepared in T-cell medium at 5 × 10^5^ cells per mL. Vybrant DyeCycle Ruby stain (1 μL) was added to the cell suspension and mixed well at a final concentration of 5 μM, followed by incubation at 37 °C for 30 min in the dark. The cells were then analyzed without washing on a flow cytometer, using fluorescence excitation and emission maxima of 638 and 686 nm, respectively.

### Imaging and single cell tracing

T cells and Dynabeads were imaged using an IX51 fluorescence microscope (Olympus, Japan) and an SLR camera at 40× magnification, on Invitrogen Countess cell counting slides (Thermo Fisher Scientific). The distance that the cells moved towards the beads was recorded and traced between the edges of the objects using Image J. Distances were then fitted to a cubic spline for extremely nonlinear relationships. Polar histogram analysis was performed using MATLAB.

### Primary human T cell electroporation

The 3 kbp DNA plasmid encoding green fluorescent protein (GFP) (Lonza) was used to assess transfection efficiency. Electroporation was performed on the human T cells on day 3 of expansion, according to manufacturer’s protocol, using cuvettes (100 μL), resulting in 1 × 10^6^ cells per electroporation with 1 μg of the plasmid. The mixture was then gently transferred to T cell-medium and cultured at 37 °C in a humidified atmosphere. Flow cytometry was performed 24 h after electroporation, to assess green fluorescence expression.

### Statistical analysis

All experiments were conducted as at least three replicates. Statistical analyses were conducted using Graphpad Prism v. 9 and data were presented as mean ± s.d.. The coefficient of determination (R^2^) was used to indicate goodness of fit. Unless otherwise stated, groups were compared using two-tail t tests.

## Acknowledgments

The authors would like to thank Dr. Polly Fordyce for helpful discussions and advice on experimental design. The authors thank the training staff of the UCI medical center for their help in obtaining blood samples from consenting breast cancer patients.

## Funding

National Science Foundation and the industrial members of the Center for Advanced Design and Manufacturing of Integrated Microfluidics (NSF I/UCRC award number IIP 1841509); American Cancer Society Institutional Research Grant (IRG-19-145-16).

## Author contributions

R. J. initiated the study, designed and implemented the experiments and analysis, and wrote the manuscript.

Y. C. prepared samples, performed electroporation and contributed to data analysis.

R. P. prepared patient blood samples.

A. A. provided guidance and contributed to data analysis.

A. P. L. supervised the research and edited the manuscripts.

## Competing interests

The authors declare no competing financial interests.

## Data and materials availability

All data are available in the main text or the supplementary materials.

## Supplementary Materials for

**This PDF file includes:**

Supplementary Figs S1 to S14

Supplementary Table 1

Supplementary Video 1

**Supplementary Figure S1.**
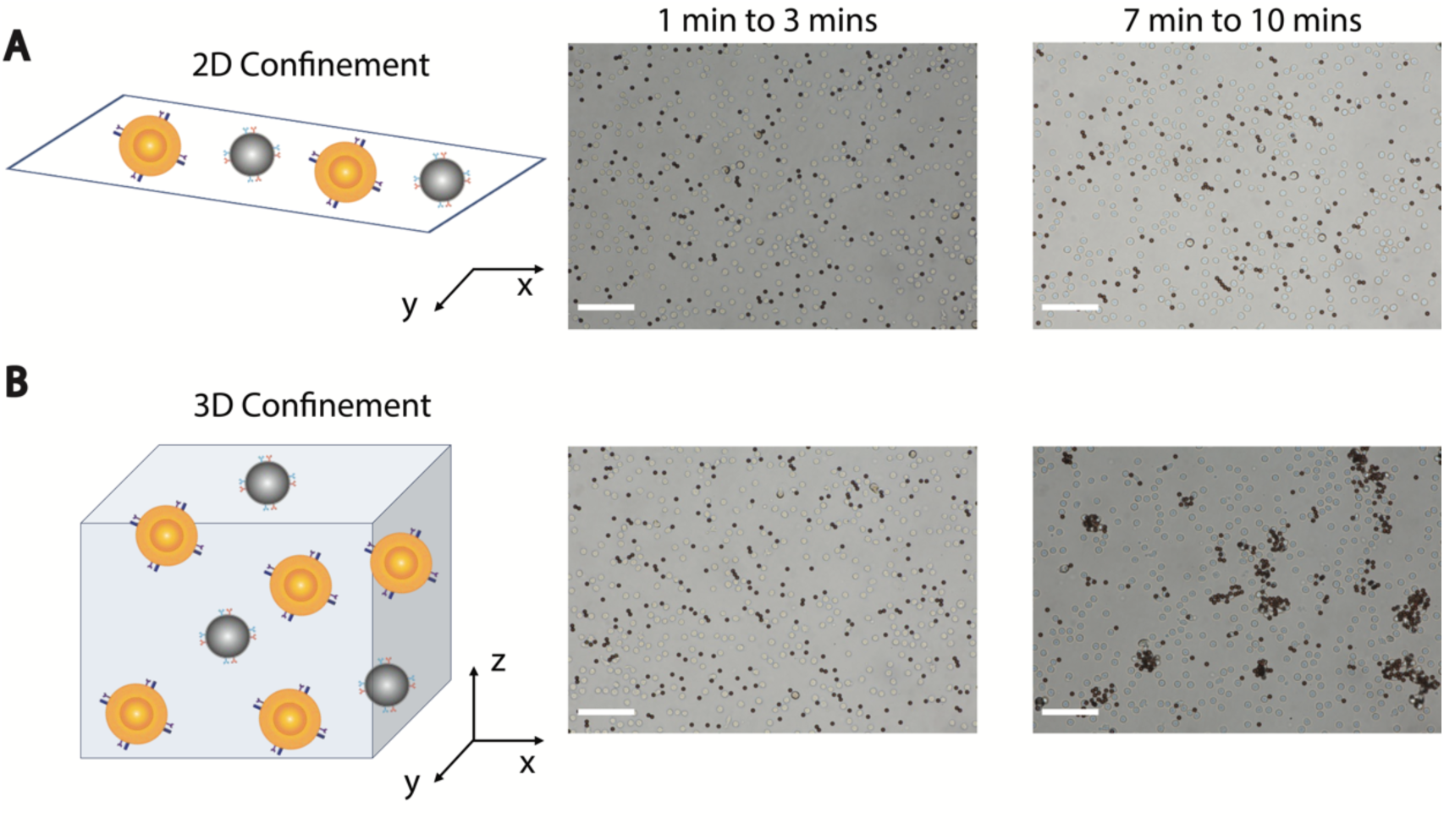
3D and 2D confinement interaction phenotypes observation. 3D confinement vs 2D confinement interaction conditions, at 100 million particles per mL (cell:bead ratio = 1:1). A. 2D confinement for the 1–3 and 7–10 min incubation periods. B. 3D confinement for the same periods. Scale bar: 50 μm.

**Supplementary Figure S2.**
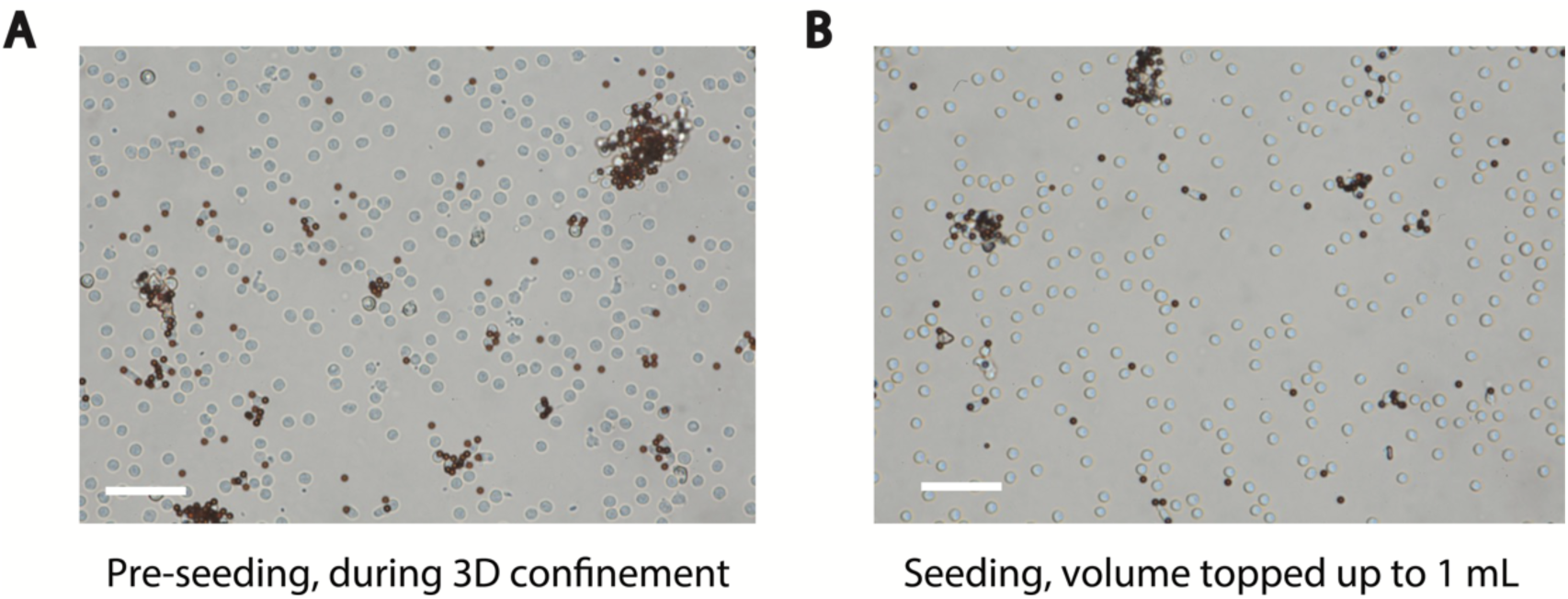
Pre-seeding vs seeding of T cell–Dynabead skeletons. A. Bright field images of cell–bead skeletons prior to volume filling and seeding, showing the more visible cell-bead skeletons. B. Bright field images of cell–bead skeletons after volume filling. Less visible cell–bead skeletons and unstable cell–bead junctions have been removed. Scale bar: 50 μm.

**Supplementary Figure S3.**
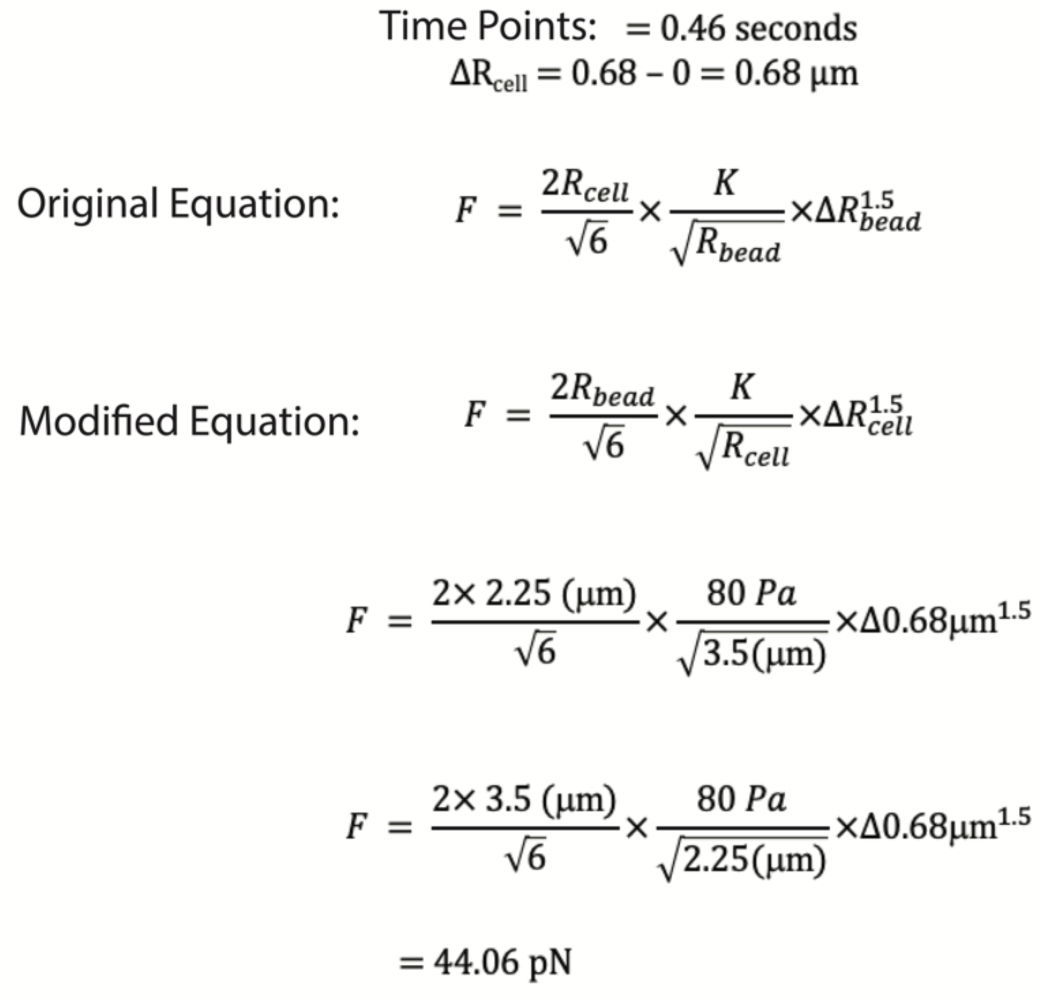
Force derivation for human T cells. The time point when the pushing or pulling movement has stopped is selected. The changes in distance are taken to represent changes in the T cell membranes.

**Supplementary Table S1.** Distance that the Dynabead was pushed by the T cells. These times and distances are used in the example in Supplementary Figure S3.

| Time (seconds) | Distance (μm) |
| --- | --- |
| 0 | 0 |
| 0.1 | 0.22 |
| 0.23 | 0.37 |
| 0.3 | 0.52 |
| 0.36 | 0.58 |
| 0.46 | 0.68 |
| 0.63 | 0.47 |
| 0.73 | 0.36 |

**Supplementary Figure S4.**
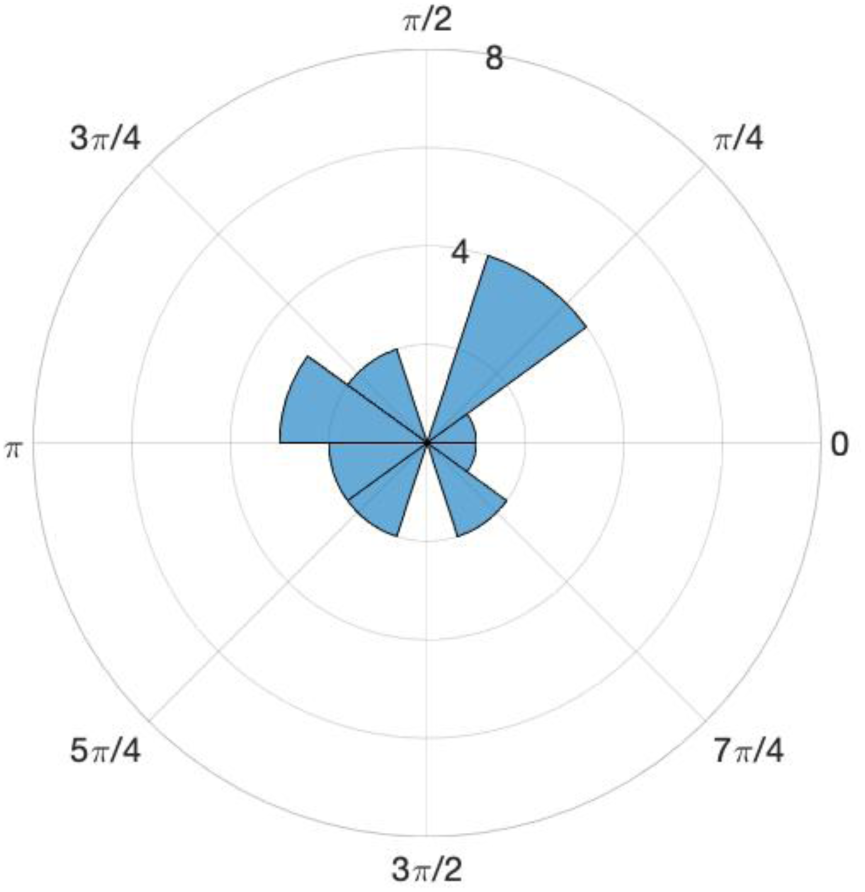
Rotational analysis of unbonded T cells and beads. The motion of T cells and beads were tracked for 5 mins.

**Supplementary Figure S5.**
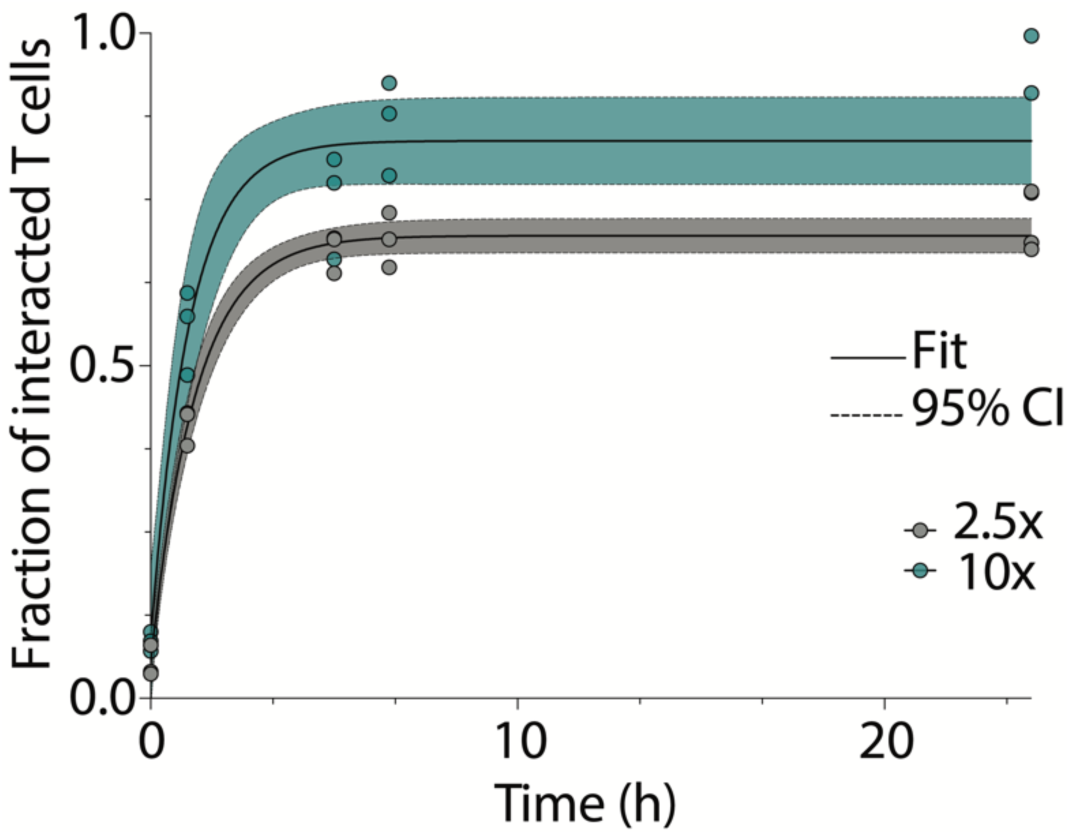
3D confinement under different concentrations. Fitted rate constant of bead–to–T cell joining in free solution (5 million and 10 million particles per mL). Black curve: least-squares fit. Black dashed lines: confidence intervals representing two standard deviations.

**Supplementary Figure S6.**
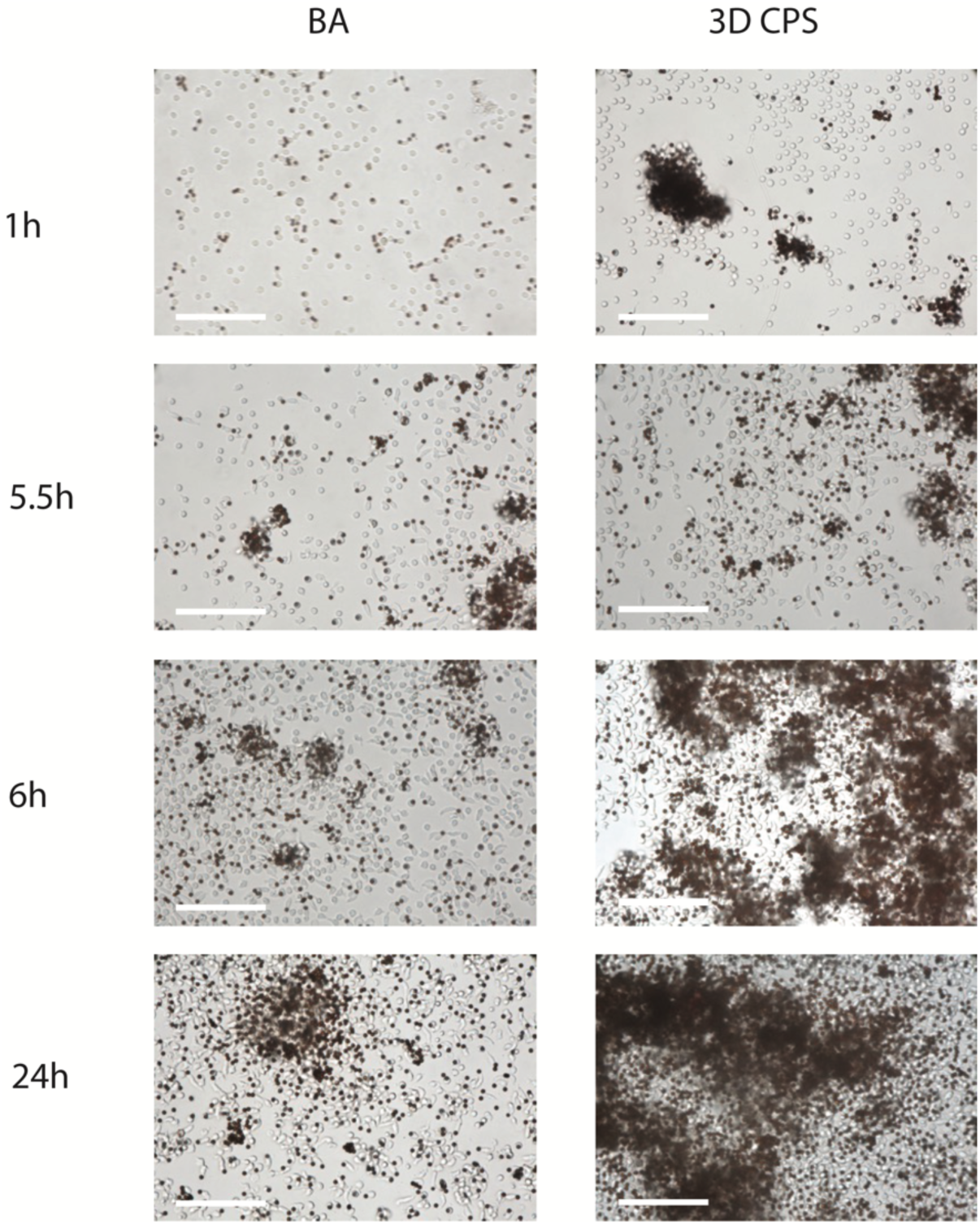
Additional activation T cell example images. Additional bulk and 3D confinement groups activation kinetics over the 24 h period. Scale bar: 50 μm.

**Supplementary Figure S7.**
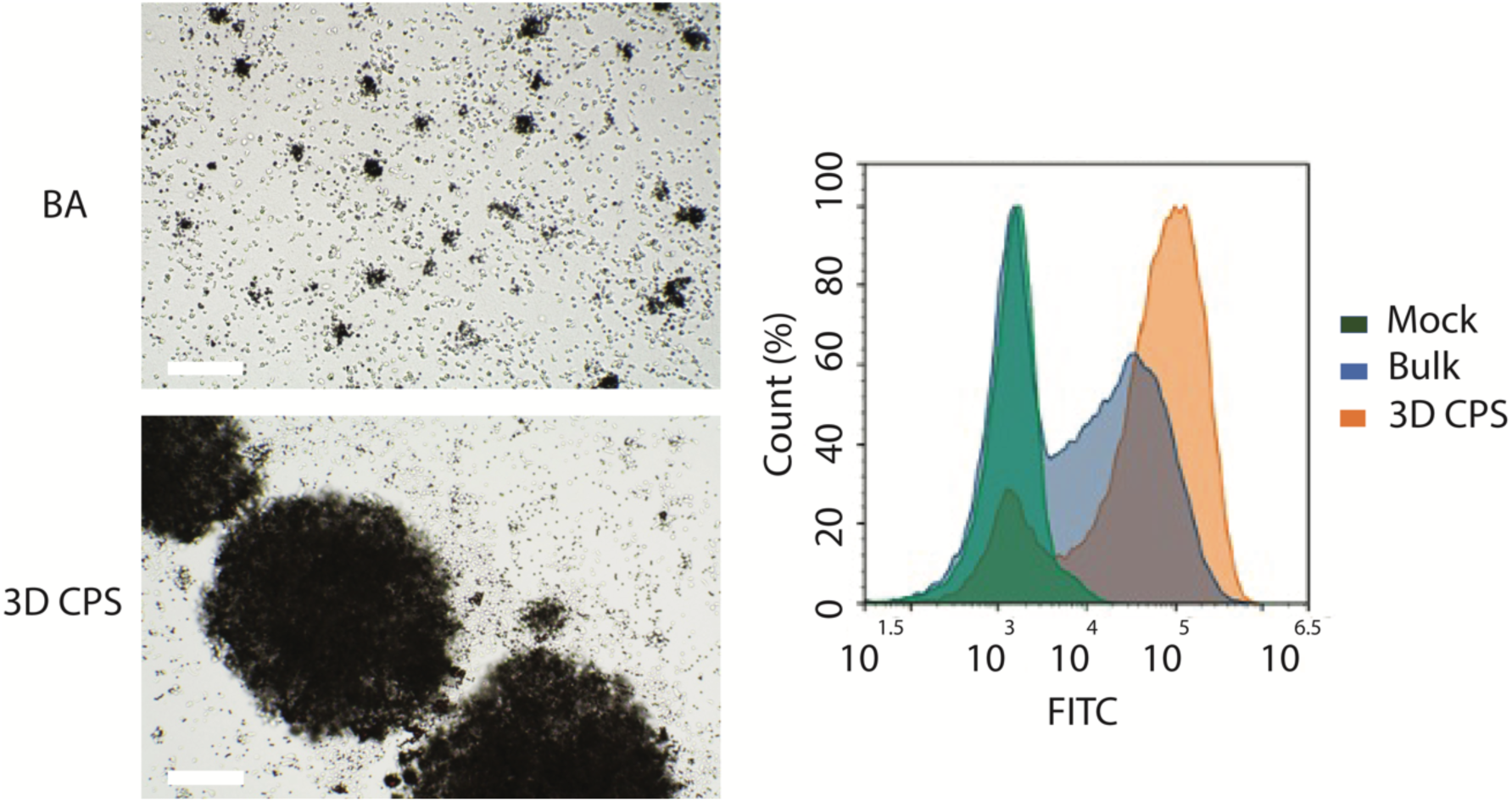
Additional activation analysis. Additional activation phenotypes for the two groups (bulk activation vs 3D CPS) on day one. Flow cytometry was conducted on day one after imaging. Scale bar: 100 μm.

**Supplementary Figure S8.**
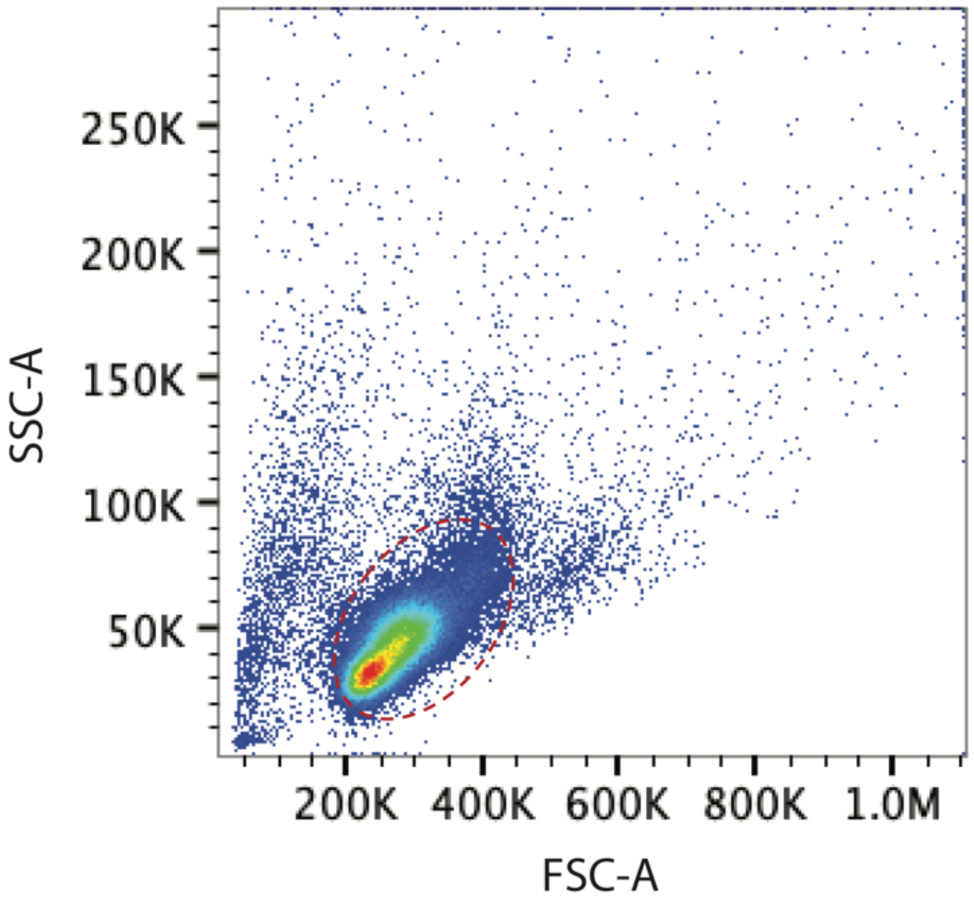
Flow cytometry gating strategy. Red circle: T cell population gated for our analysis.

**Supplementary Figure S9.**
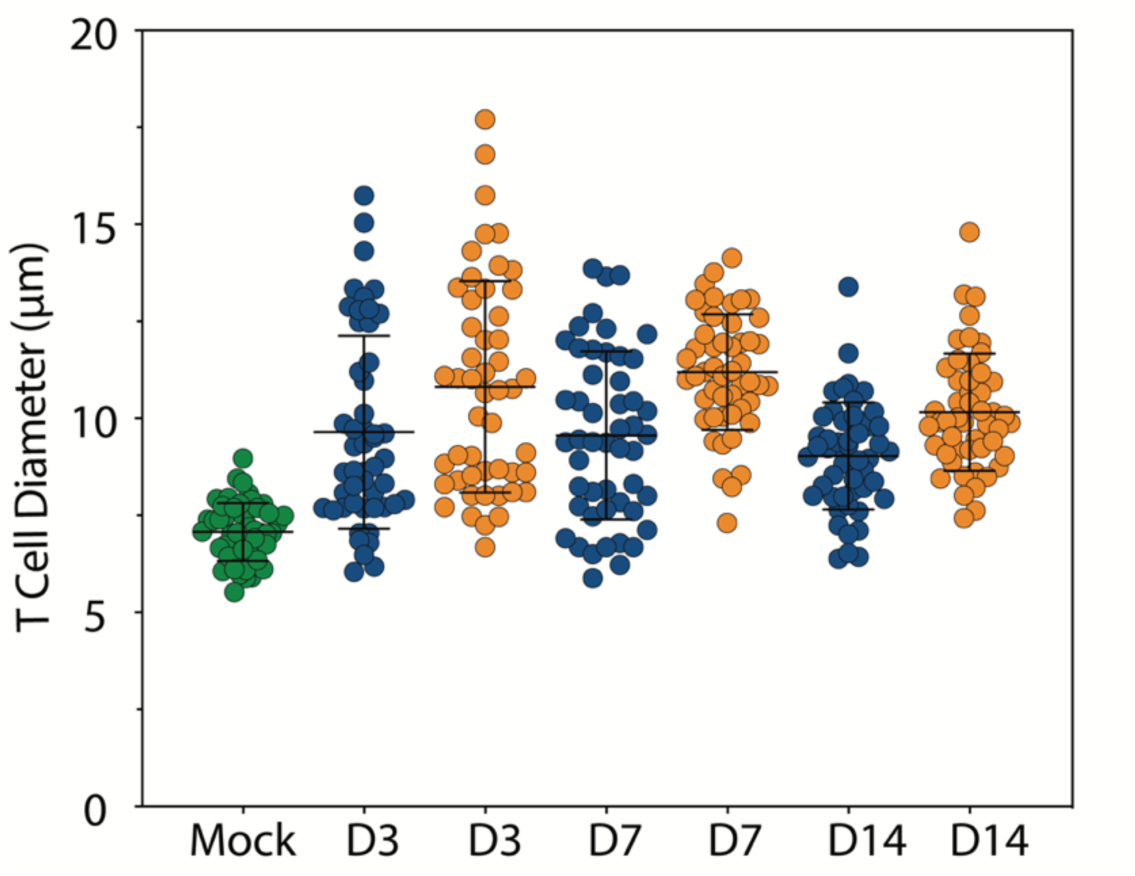
Primary T cell size distribution. Primary T cell size distribution over the two-week clonal expansion period. Green: T cells without activating molecules; blue: bulk activation group; orange: 3D confinement group.

**Supplementary Fig. S10.**
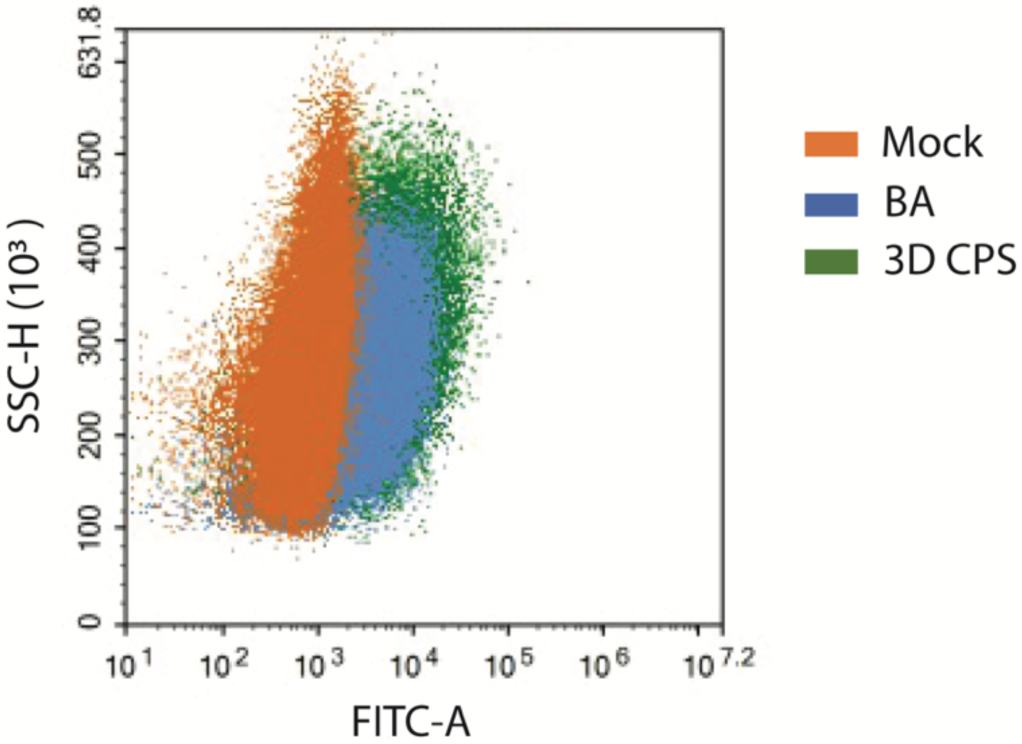
Exhaustion marker analysis. PD-1 expression analysis via flow cytometry on day 14 of clonal expansion.

**Supplementary Figure S11.**
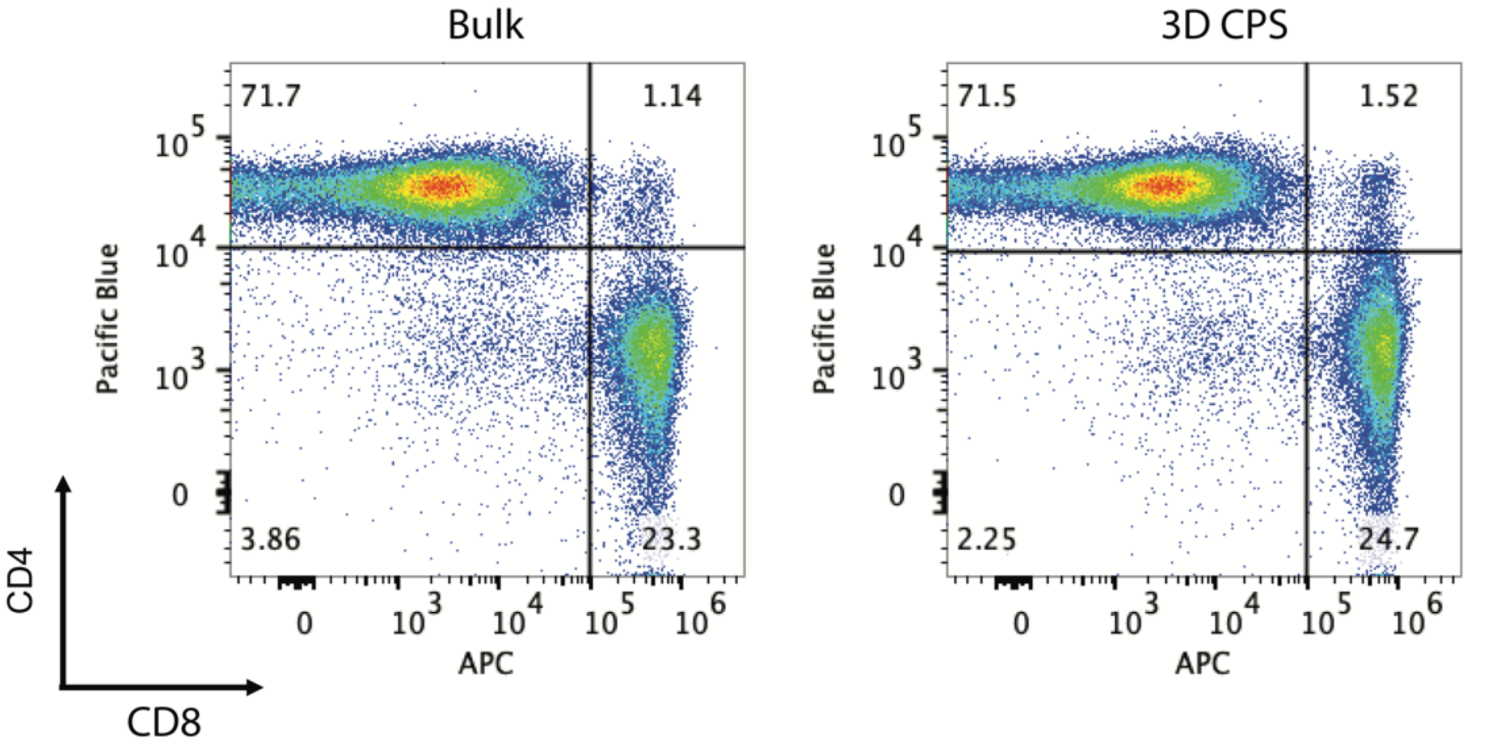
Ratio of CD4 T cells to CD8 T cells. CD4:CD8 ratio on day 14 of T cell clonal expansion. CD8 was labeled with APC labeled and CD4 was labeled with pacific blue.

**Supplementary Figure S12.**
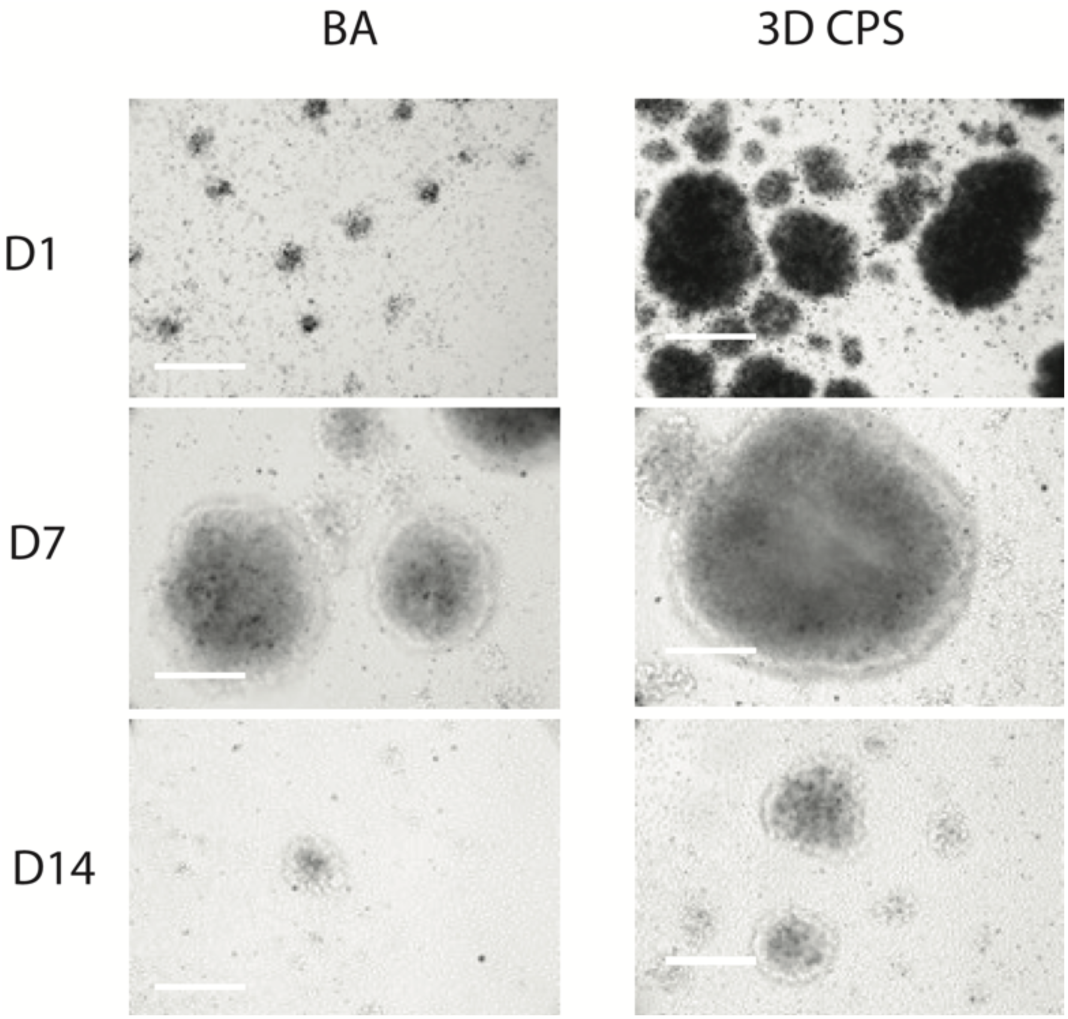
Additional T cell expansion images. Additional human T cell expansion images on days 1, 7, and 14. Scale bar: 100 μm.

**Supplementary Figure S13.**
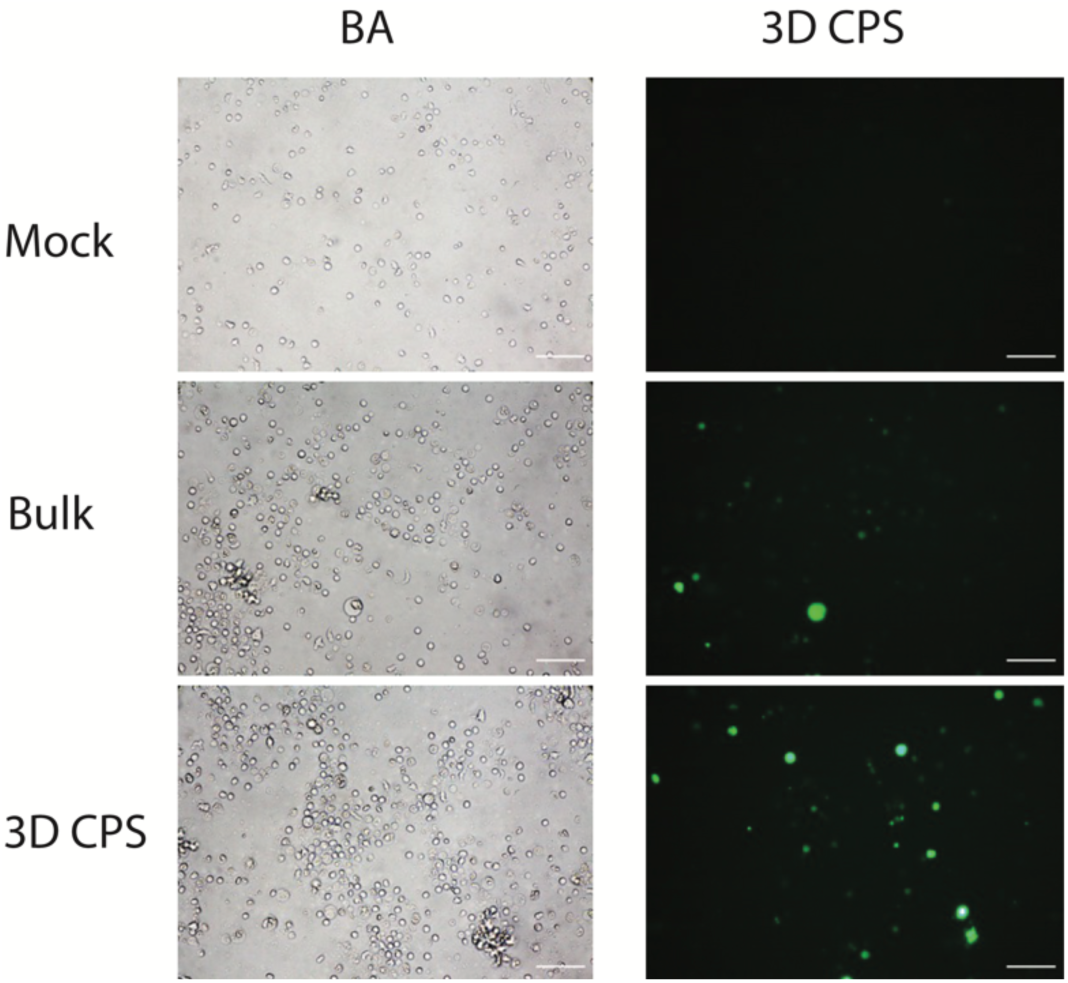
Additional primary T cell transfection images. Plasmid electroporation for the three groups (mock, bulk activation, and 3D confinement). Bright field and fluorescent images of transfected primary T cells. Scale bar: 50 μm.

**Supplementary Figure S14.**
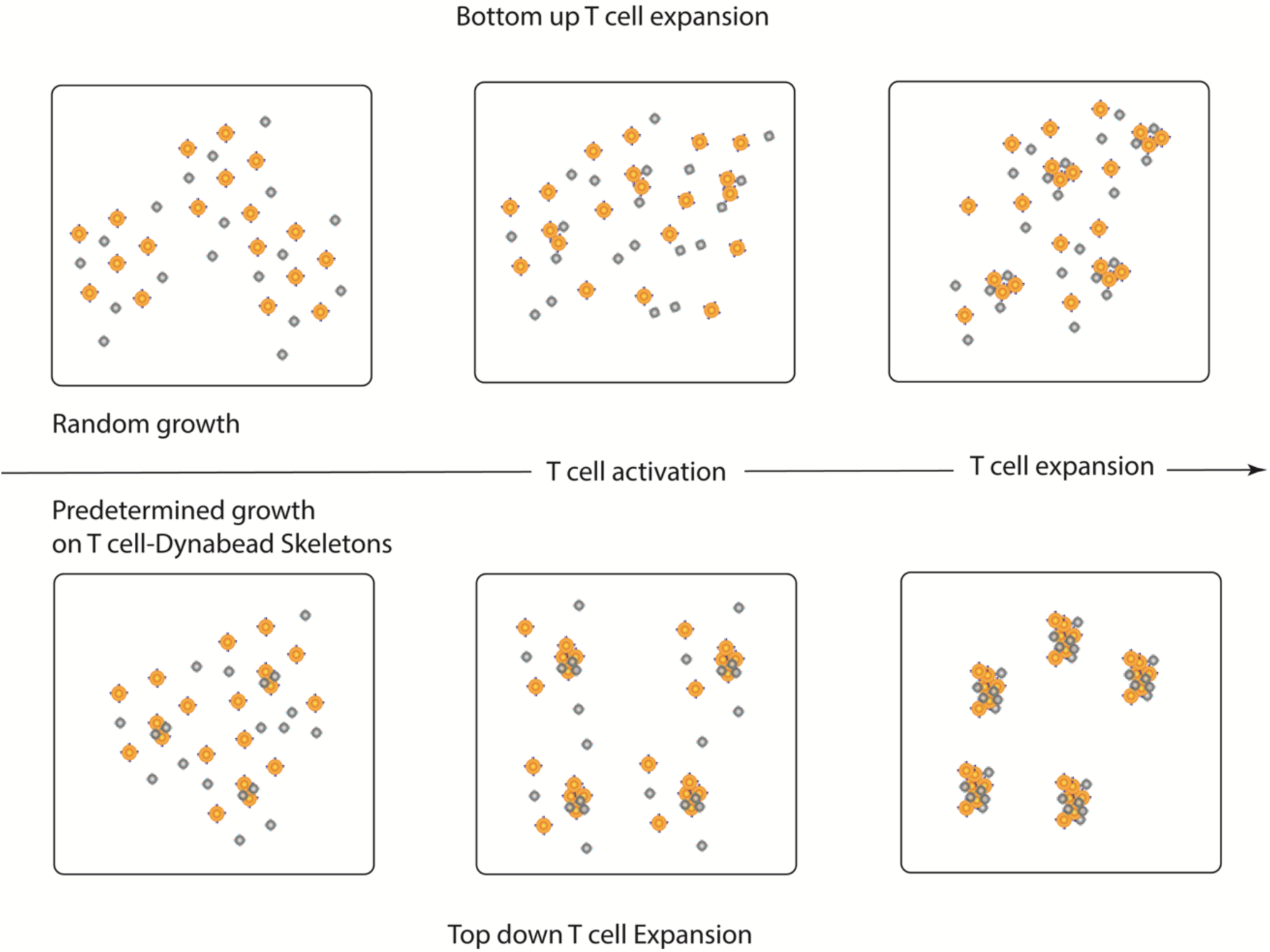
Bottom up T cell expansion and top down T cell expansion workflow comparison. Top down T cell expansion workflow offers precise and ordered control from T cell-Dynabead skeletons and overcomes stochasticity from bottom up T cell expansion approach.

## Notes

### Competing Interest Statement

The authors have declared no competing interest.

